# Single camera estimation of microswimmer depth with a convolutional network

**DOI:** 10.1101/2025.05.09.653055

**Authors:** Ali Hosseini, Célia Fosse, Maya Awada, Marcel Stimberg, Romain Brette

**Author notes:** Corresponding author: Romain Brette.

## Abstract

A number of techniques have been developed to measure the three-dimensional trajectories of protists, which require special experimental setups, such as a pair of orthogonal cameras. On the other hand, machine learning techniques have been used to estimate the vertical position of spherical particles from the defocus pattern, but they require the acquisition of a labeled dataset with finely spaced vertical positions. Here we describe a simple way to make a dataset of *Paramecium* images labeled with vertical position from a single 5 minutes movie, based on a tilted slide setup. We used this dataset to train a simple convolutional network to estimate the vertical position of *Paramecium* from conventional bright field images. As an application, we show that this technique has sufficient accuracy to study the surface following behavior of *Paramecium* (thigmotaxis).

## Introduction

The natural environment of swimming microorganisms is three-dimensional. Protists such as the common freshwater ciliate *Paramecium* often swim upward towards the surface (Häder and Hemmersbach, 2018; Roberts, 2010), but can also dive down (Applewhite and Gardner, 1973), hover above surfaces or land on them (Iwatsuki and Hirano, 1995; Ohmura et al., 2018), and explore the full volume of their environment (Brette, 2021; Jennings, 1906). Yet, due to technical constraints, observation is generally restricted to a two-dimensional projection.

A number of three-dimensional observation techniques have been designed using special equipment. Drescher et al. (2009) reconstruct trajectories from two orthogonal cameras. This is precise, although with a limited observable volume. Other techniques use custom microscopes, such as the tPOT microscope, which uses a principle akin to stereomicroscopy (Marumo et al., 2021; Yajima et al., 2008). Others use a multicolored light sheet (Matsushita et al., 2004; McGregor et al., 2008), a custom laser-based feedback system (Peters et al., 1998), a confocal microscope (Dinsmore et al., 2001; Rabut and Ellenberg, 2004) or holographic microscopes (Daloglu et al., 2018; Memmolo et al., 2015; Su et al., 2013). Several authors have developed three-dimensional tracking using a motorized stage or microscope (Darnige et al., 2017; Hasegawa et al., 2008), but this is necessarily limited to the observation of a single cell, and requires careful synchronization of camera and motor control.

All these techniques rely on special equipment that is not usually available in labs. Another possibility to estimate the 3D position of particles from conventional microscope images is to infer the vertical position from the defocus pattern. This approach is called *defocus particle tracking* (Barnkob et al., 2021). It is based on correlation with images of known position (Taute et al., 2015) or trained neural networks (König et al., 2020; Newby et al., 2018; Ratz et al., 2023; Zhang et al., 2022). In either case, it requires the prior acquisition of a stack of images at finely spaced vertical positions.

Defocus particle tracking has been applied to spherical particles, typically for particle tracking velocimetry, but not to protists. The additional difficulty is that protists have more variable and non-symmetrical shapes, and they swim. This makes both acquisition and training more challenging. Here we train a convolutional network to estimate the vertical position of *Paramecium* from conventional bright field microscope images, using a simple trick to acquire a labeled dataset. The trick consists in placing swimming paramecia between two tilted glass slides, so that the vertical position *z* correlates with the *x* coordinate, which is used as a proxy for *z* during training. This allows training the network with a single continuous 5 minutes movie. We show that the trained model can then estimate the vertical position of cells on a horizontal slide. We demonstrate the applicability of the method to the study of thigmotaxis (attraction to surfaces).

## Methodology

### Experimental procedures

Specimens of *Paramecium tetraurelia* were obtained from Éric Meyer, Institut de Biologie, Ecole Normale Supérieure, Paris, France. Cells were grown at room temperature (about 22°C) with a light-dark cycle (12 hours) in a wheat grass powder (Pines International, Lawrence, KS) infusion medium bacterized with *Klebsiella pneumoniae* the day before feeding the cells and supplemented with 0.8 µg/ml β-sitosterol.

For the training datasets, we mixed stationary and log-phase cells, which have different sizes and shapes (Fig. 1). Cells were washed for at least 30 minutes in a salt solution that mimicked the ionic composition and pH of the culture medium. The solution contained 0.8 mM KCl, 0.1 mM CaCl_2_, and 0.64 mM NaCl, buffered with 1.4 mM NaH_2_PO_4_·H_2_O and 4.2mM Na_2_HPO_4_·2H_2_O, with the pH adjusted to 7.5 using NaOH, resulting in a total Na concentration of approximately 11.5 mM. To account for the difference in cell densities – around 1000 cells/mL for log-phase cultures and over 4000 cells/mL for stationary-phase cultures – a 4:1 volume ratio of log-phase to stationary-phase cells was used. All chemicals were acquired from Sigma-Aldrich.

**Figure 1.**
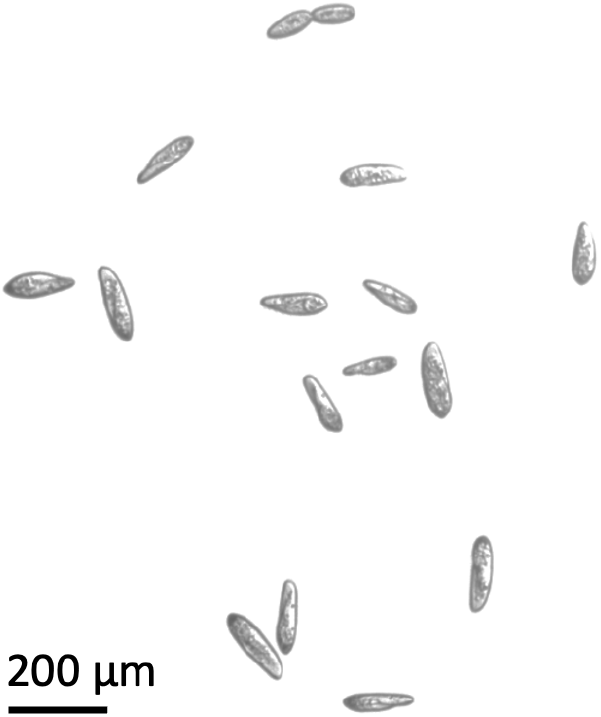
Mixture of log and stationary cells. A dividing cell can be seen on top.

We trained and tested the models on a motorized upright microscope with a 4x objective (Olympus, NA 0.10, FN 22) and a 6 MP camera (Lumenera fnfinity 3-6UR; 8 bits depth, 2752 x 2192 pixels), with bright field illumination. Pixel size was 1.78 µm.

To make a test dataset with controlled depth, a drop of 40 µl of cell mixture was put on a plain glass slide (Premium Plain Slides, Epredia, 2950WX-003 25x75x1 mm, USA) and gently covered by a square glass coverslip (20 x 20 mm). This flattened the drop on the surface and limited the movement of the cells to virtually a 2D plane. Movies were taken at depth 0 to 500 µm in steps of 100 µm, using a motorized stage (Luigs and Neumann, D-40880, Germany).

To make datasets with a tilted slide, a drop of the cell mixture was placed on a plain glass slide and sandwiched by placing a second glass slide over it with a fixed 260 µm distance (using tape). The slide sandwich was then inclined such that the distance between the maximum and minimum heights within the field of view was approximately 1 mm, corresponding to an angle of 9.4 degrees relative to the horizontal plane (Fig. 2A). The microscope image was then focused on top. In this way, defocus increases from right to left (Fig. 2B).

**Figure 2.**
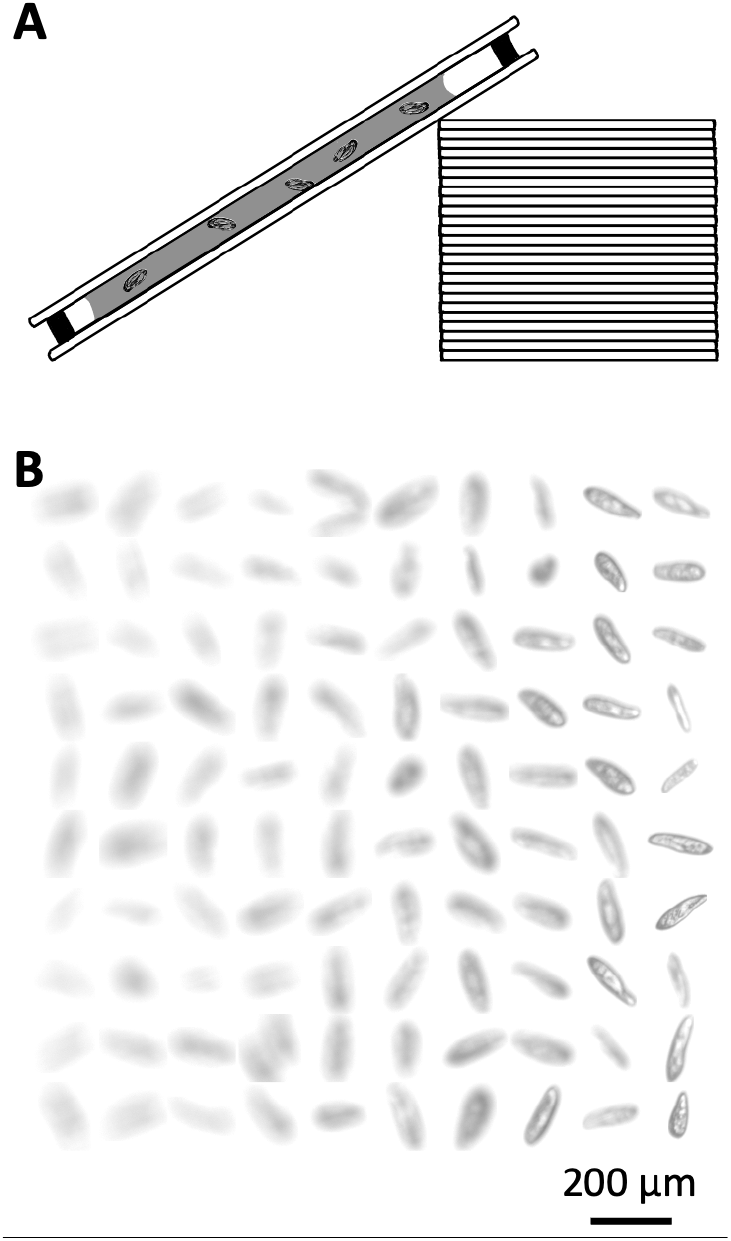
Recording on a tilted slide. A, Glass slide sandwich resting on a pile of slides. B, Sample cell images ordered from left to right by horizontal position on the slide.

For the application to thigmotaxis, a drop of culture medium with stationary cells was placed between two horizontal glass slides, spaced by 520 µm (using tape), and the image was focused on top.

### Data acquisition

Images were acquired at a fixed frame rate (2, 10 or 20 Hz). Light intensity, exposure and gain were varied to test for robustness. The background was calculated over a few seconds (mean of images) then subtracted online, and the resulting image was thresholded then saved with lossless compression. This ensured manageable data size for storage.

We then extracted images of individual cells. This was done with conventional tracking techniques. We used Fasttrack (Gallois and Candelier, 2021), which uses classical computer vision methods: binarization of the threshold image, selection of connected components of adequate size, and calculation of the centroid. Then a square image around the centroid was extracted, of size 96 pixels (171 µm). Squares extending beyond the image boundaries were discarded. To make the training datasets, we removed tracks smaller than 300 µm, in order to avoid still cells and errors.

Training datasets were built from 5 minutes movies at 20 Hz, totaling 6000 frames, from which about 100,000 cell images were extracted. For each image, we assigned a label corresponding to the central vertical position of the slide sandwich given the cell’s horizontal position (i.e., expected position of the cell). The test dataset with controlled depth was acquired at 2 Hz over 5 minutes. The application to thigmotaxis was done from a 3 minutes movie acquired at 10 Hz.

## Models

### Architecture

We used a simple network with two convolutional layers with kernel size 3 and two dense layers (Fig. 3). The first convolutional layer has 32 filters and is followed by a max pooling operation (size 2), and the second one has 64 filters and is followed by global average pooling. The output is flattened and fed to a dense layer with 128 output units. A final dense layer outputs a single number, which is the *z* estimate. All units have leaky rectified linear activation (leaky ReLU), except the final unit, which is linear. Thus, the model produces estimates based on local texture.

**Figure 3.**
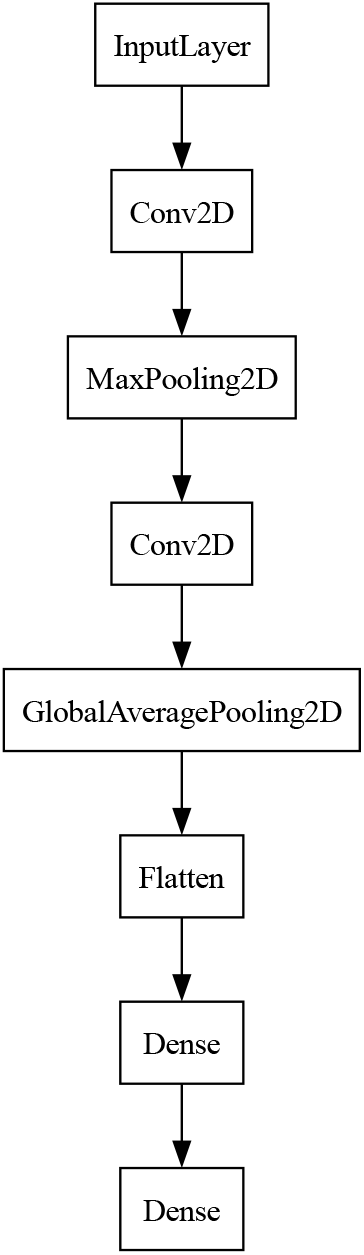
Model architecture.

### Preprocessing and augmentation

Images are inverted so that the background is zero. The final 20% of the movie is used for validation, then the frames are shuffled in both training and validation sets. In the training dataset, images are augmented with random horizontal and vertical flipping, and Gaussian noise is added, with a standard deviation taken for each image from a uniform distribution between 0 and 5 (out of a range of 0 to 255).

### Training

Models were optimized for quadratic loss (mean squared error) relative to the assigned vertical position. They were trained for up to 500 epochs on a MacBook Pro (36 MB; M3 Pro) using Adam optimizer, starting with a learning rate of 5.10^-4^, halved when the loss stalls or increases. The weights with lowest mean absolute error on the validation dataset during training were saved.

### Prediction

With the tilted slide setup, the model is trained on variations of distance in air between the microscope and the cells. Thus, when the model is used to estimate the depth of cells in water, a corrective factor of 1.33 must be applied to account for the difference in refraction index between air and water. This factor is applied in the thigmotaxis application (Fig. 7), but not in the various tests, which are done with variations of distance in air.

## Results

### Training and testing the model

To make a dataset representative of the diversity of shapes, we mix cells in logarithmic phase (large cells) with cells in stationary phase (smaller starved cells), as illustrated on Figure 1. The apparent shape of cells also changes depending on their angle relative to the horizontal plane. In order to allow for diverse cell orientations, we place a drop of cells between two glass slides spaced by 260 µm, which is about twice the cell length (120 µm). The slide sandwich is then tilted by raising one end with a stack of slides (Fig. 2A), to an angle of about 10°. In this way, the vertical position of cells (*z*) varies systematically with the horizontal position (*x*), over a range of about 800 µm (Fig. 2B).

We then take a 5 minutes movie. Images are preprocessed by removing the background and thresholding, then small images of individual cells are extracted. The model is then trained to predict the cell’s expected vertical position, as inferred from the cell’s horizontal position. After training, the model obtains a mean absolute error of 48 µm on the validation set (the last minute of the movie). Note that with our tilted slide design, it is the distance in air, not in water, that varies. As the refraction index of air and water differ, this must be corrected. A precision of 48 µm in air is optically equivalent to 64 µm in water (48 µm times 1.33).

Note that part of this estimation error comes from the fact that, since the two glass slides are spaced by 260 µm, there is some variability in the actual vertical position of the cell at a given horizontal position along the slide, as can be observed in Figure 2B. We can estimate this contribution as follows. Cells are about 35 µm wide, plus 10 µm for cilia. This means that the maximum distance between two cell centers is 205 µm. If cells are uniformly distributed between the two slides, then the average distance of a cell center to the center of the slide sandwich is 51 µm. In summary, the precision of the model’s estimation is comparable to the variability of cell position within the slide sandwich.

When we look at the model’s estimates in detail as a function of *z* (Fig. 4), we notice a bias on the boundaries of the training set, at minimum and maximum depth. This is to be expected because a good estimator would take constraints of the dataset into account, and therefore output estimates within these bounds. This means that when *z* is at maximum depth, the estimate will always be above it, and therefore biased. This bias can be mitigated on the defocused end (low z) by including a large enough range of vertical positions in the training set, but this cannot be easily done on the focused end (z near 0). Thus, estimates near focus tend to be biased.

**Figure 4.**
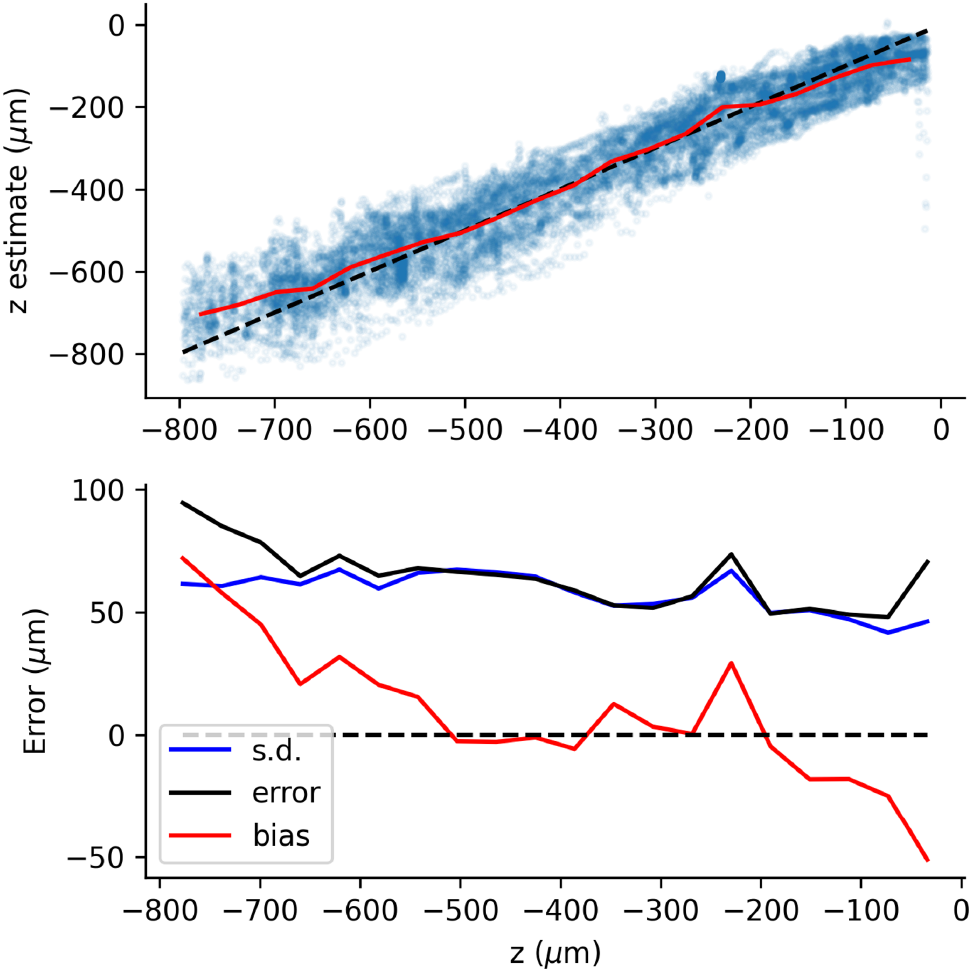
Estimation results after training. Top: Estimates of z after training, as a function of expected z (blue line: perfect estimation; red: mean estimate as a function of expected z). Bottom: root mean square error (black), standard deviation (black) and bias (red), vs. expected z.

One practical challenge, which was not assessed in previous work on defocus particle tracking, is that recording parameters, such as exposure time, light intensity and camera gain, can vary across recordings. One obvious solution is to train the model for each recording condition. To make it more practical, we trained the model by augmenting the dataset with noise, so as to make the model more robust to varying recordings conditions. Figure 5A shows estimation results on a tilted slide sandwich, with different recording parameters compared to the training dataset (higher gain, higher light intensity and lower exposure time). These results are similar to the validation set (error of 46 µm vs. 48 µm). When noise is not added during training, the model is not robust, as shown on Figure 5B. SpeciIically, in the test slide, which is noisier (higher gain), the estimated position of defocused cells is biased towards more focused cells. As a result, the linear regression between the estimate and the actual *z* has slope 1.34 (vs. 0.96 when trained with noise).

**Figure 5.**
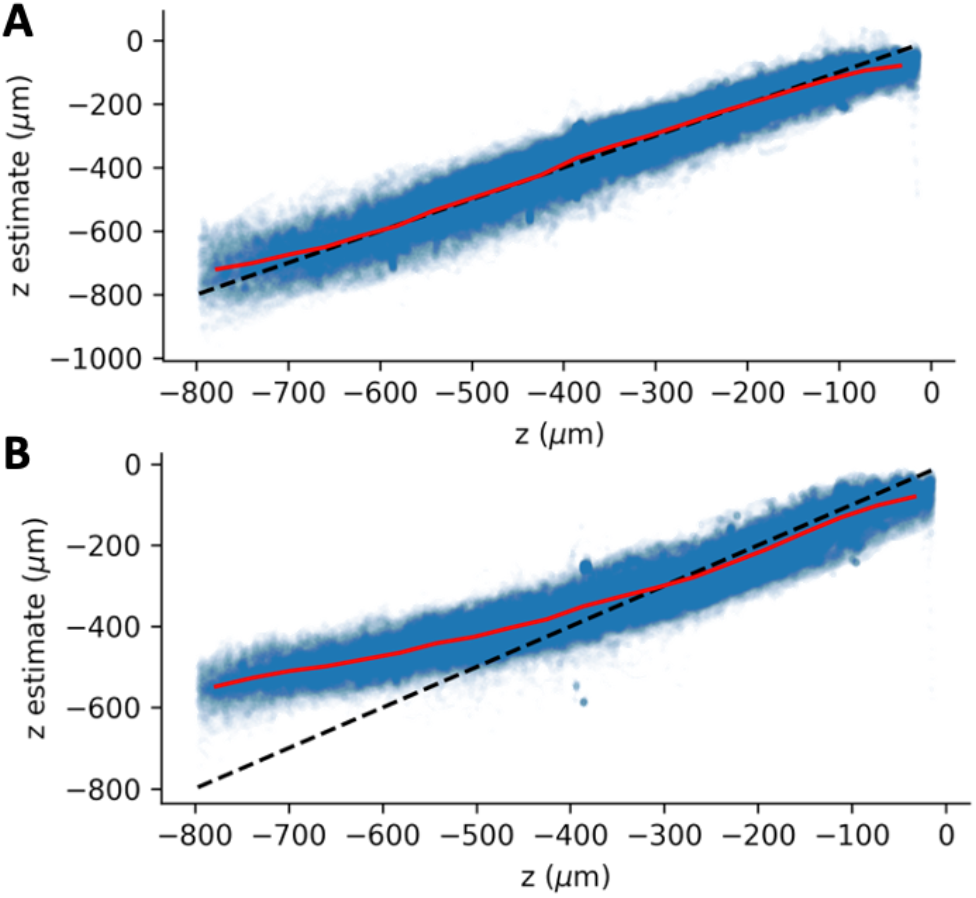
Estimation results with different recording settings. A, With noise added during training. B, Without noise augmentation (same legend as Fig. 4A).

We then measured the precision of estimates with ground truth measurements of *z*. To this end, we placed a thin film with cells on a glass slide, and took movies with the microscope focus positioned 0 to 500 µm above the cells, in steps of 100 µm, using a motorized microscope. The average absolute error was 46 µm, similar to the error obtained on tilted slides, and the regression slope was 0.99 (Fig. 6A).

**Figure 6.**
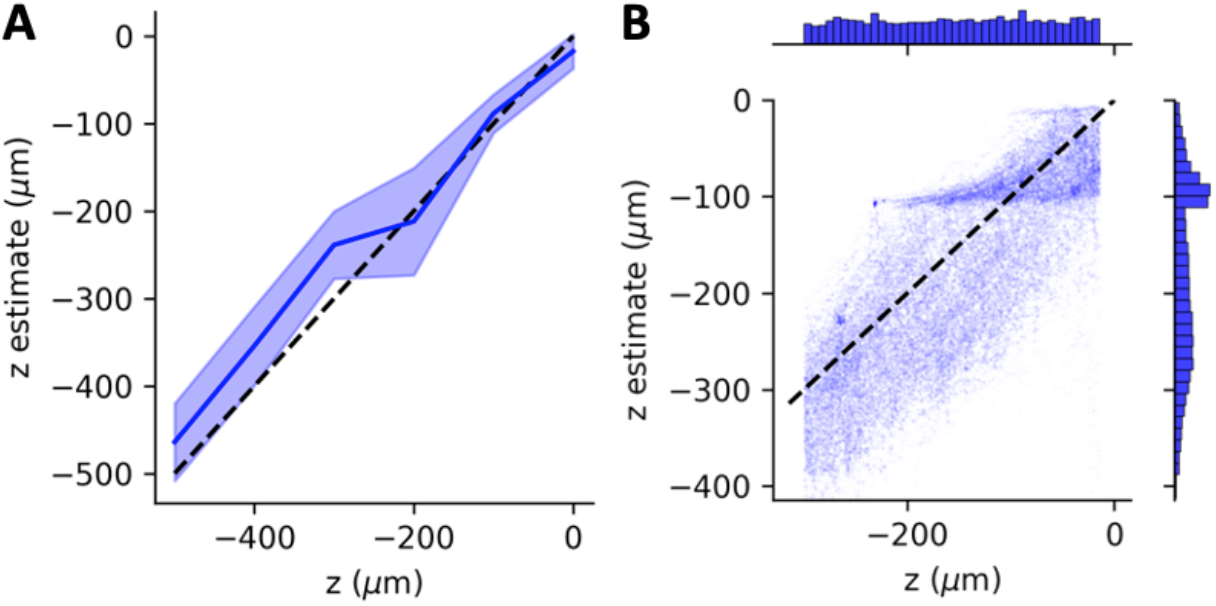
Testing the trained model on recordings with controlled depth. A, Model estimate vs. true vertical position (dashed line: identity relation; shaded area: mean estimate ± mean absolute error). B, Training on Pixed depths and testing on tilted slide produces a ladder effect.

If instead, the model is trained on this discrete set of recordings, we obtain an error of 27 µm, but of course this is a much simpler task since the set of possible positions is very small (6 possibilities). In addition, in this dataset, cells are constrained to be horizontal and therefore do not reflect the full diversity of cell appearances. When we test this model on a tilted slide, the error is similar to previous results (46 µm), but as can be seen on Figure 6B, there is a ladder effect: near focus, there is a peak in the distribution of estimates near -100 µm, which is one of the positions of the discrete dataset.

### Application

We now show an application of this estimation model to the study of thigmotaxis. Thigmotaxis is the tendency of some microorganisms to follow surfaces or attach to them. This is a major part of the life of *Paramecium* in its natural habitat (Jennings, 1906), but it has not been the object of many studies, possibly for technical reasons. We simply placed a drop of culture medium with stationary cells (i.e., starved) between two glass slides, spaced by 520 µm, and took a 3 minutes movie.

Figure 7A shows the estimated position of cells on part of a frame (full annotated movie: Supplementary Movie 1). Note that the water refraction index is taken into account by multiplying the model estimate by 1.33. One of the cells is dividing. Figure 7B shows the speed and vertical position of all cells measured in all frames. We can see that most cells are concentrated near the two surfaces. Many are swimming and a few cells are attached, both on the top and the bottom surfaces. Regarding the *z* estimates, we notice that the attached cells are estimated away from the surfaces by ∼100 µm or more. First, we expect the estimate to correspond to the cell center. Given that cell width is on average 34 µm (Nagel and Machemer, 2000) and cilia are 10 µm long, this means the center of an attached cell is at least 27 µm away from the surface. Second, cells tend to attach with their anterior end, slightly tilted, which moves away the cell center from the surface (e.g. about 40 µm for a 30° angle). The rest might be accounted for by the precision error and biases, especially near focus, as we have seen previously. Looking more closely at the attached cells (zero speed), we can also see some positive estimates, which must be erroneous. Looking at the annotated movie, this appears to be due to occlusions due to background removal, when the cell was attached during the calculation of the background and later moved. Finally, we also notice some unusually low estimates for attached cells, with a rather broad distribution (down to -200 µm). Looking again at the movie, this appears to be due to the dividing cell shown on Figure 7A, which is initially horizontal and later turns on the anterior end of one of the daughter cells, so that the dividing cell is almost vertical, and this is accompanied by a decrease in the *z* estimate. Thus, these low values of *z* actually seem genuine.

**Figure 7.**
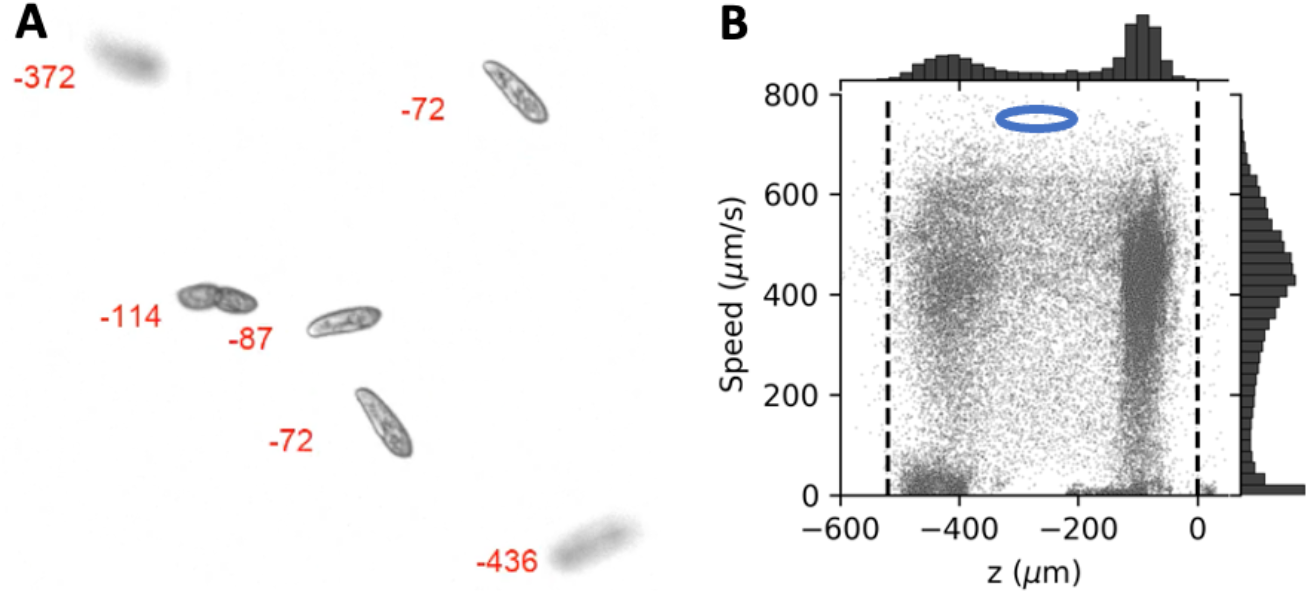
Application to the study of thigmotaxis (A, B). A, Estimated position in red (µm). B, Speed vs. z for all measured points, with marginal distributions. The blue ellipse shows the typical size of a cell for scale (120 µm long, 30 µm wide). Dashed lines: slides.

In summary, despite the limitations of the method, it seems to be able to provide useful information in the context of thigmotaxis.

## Discussion

We have introduced a simple method to train a convolutional network to estimate the vertical position of paramecia from conventional microscopy images, using tilted glass slides. With a 4x objective, we obtained a precision of about 60 µm. Augmenting the training dataset with noise improves the robustness with respect to changes in recording conditions (light intensity, focus, gain). The method can be used for example to study thigmotaxis.

Precision in the vertical dimension remains more than an order of magnitude worse than in the horizontal dimensions (pixel size was about 2 µm). This is not surprising, given that cells are about 35 µm wide and that, in contrast with the small spherical particles typically used in defocus particle tracking, they come in a variety of shapes and orientation rather than a stereotypical pattern. Nonetheless, this means that this estimation method remains inferior in accuracy to direct estimation of vertical position with an orthogonal camera, although it allows for a wider field of view. Therefore, the method does not replace dedicated equipment for accurate three-dimensional tracking, but it is useful to augment the information obtained with conventional microscopy equipment.

## Code availability

Code for the depth estimator can be found at https://github.com/romainbrette/3Dtracking. Code for tracking and background removal can be found at https://github.com/mstimberg/online_bg_removal.

## Acknowledgments

This work was supported by Agence Nationale de la Recherche (ANR-20-CE30-0025-01, ANR-21-CE16-0013-02 and ANR-23-CE16-0020-02).

